# Brain-wide human oscillatory LFP activity during visual working memory

**DOI:** 10.1101/2023.09.06.556554

**Authors:** Balbir Singh, Zhengyang Wang, Leen M. Madiah, S. Elizabeth Gatti, Jenna N. Fulton, Graham W. Johnson, Rui Li, Benoit M. Dawant, Dario J. Englot, Sarah K. Bick, Shawniqua Williams Roberson, Christos Constantinidis

**Affiliations:** Department of Biomedical Engineering, Vanderbilt University; Neuroscience Program, Vanderbilt University; Department of Neurology, Vanderbilt University Medical Center; Department of Neurological Surgery, Vanderbilt University Medical Center; Department of Electrical and Computer Engineering, Vanderbilt University; Department of Ophthalmology and Visual Sciences, Vanderbilt University Medical Center

**Author notes:** Lead Contact / Corresponding Author.

**Keywords:** LFP, intracranial recordings, working memory

## Abstract

Oscillatory activity is thought to be a marker of cognitive processes, although its role and distribution across the brain during working memory has been a matter of debate. To understand how oscillatory activity differentiates tasks and brain areas in humans, we recorded local field potentials (LFPs) in 12 adults as they performed visual-spatial and shape-matching memory tasks. Tasks were designed to engage working memory processes at a range of delay intervals between stimulus delivery and response initiation. LFPs were recorded using intracranial depth electrodes implanted to localize seizures for management of intractable epilepsy. Task-related LFP power analyses revealed an extensive network of cortical regions that were activated during the presentation of visual stimuli and during their maintenance in working memory, including occipital, parietal, temporal, insular, and prefrontal cortical areas, and subcortical structures including the amygdala and hippocampus. Across most brain areas, the appearance of a stimulus produced broadband power increase, while gamma power was evident during the delay interval of the working memory task. Notable differences between areas included that occipital cortex was characterized by elevated power in the high gamma (100-150 Hz) range during the 500 ms of visual stimulus presentation, which was less pronounced or absent in other areas. A decrease in power centered in beta frequency (16-40 Hz) was also observed after the stimulus presentation, whose magnitude differed across areas. These results reveal the interplay of oscillatory activity across a broad network, and region-specific signatures of oscillatory processes associated with visual working memory.

## INTRODUCTION

Working memory, the ability to hold and manipulate information in the mind for a short period of time, is central for many cognitive processes, including planning, problem-solving and decision-making (Baddeley, 2012; Constantinidis and Klingberg, 2016). Pathological conditions such as stroke, schizophrenia, and Alzheimer’s disease are associated with working memory impairments (Westerberg et al., 2007; Subramaniam et al., 2012; Subramaniam et al., 2021). For this reason, understanding the neural basis of working memory has been a long-standing question in cognitive neuroscience research (Buschman and Miller, 2022). Neurophysiological experiments in non-human primates have identified neuronal activation of the prefrontal cortex as central for the maintenance of working memory (Riley and Constantinidis, 2016). Persistent activity generated by the spiking of prefrontal neurons is also associated with distinct patterns of spectral power, identifiable in local field potential recordings (Singh et al., 2022; Singh et al., 2023).

However, other lines of evidence, and particularly human functional imaging studies suggest a much more widespread pattern of activation during working memory (Christophel et al., 2017). Imaging studies have been successful in using multi-variate methods to decode the remembered visual stimuli from visual cortical area voxels (Harrison and Tong, 2009). Correspondingly, whether the contents of working memory are maintained in the prefrontal or in the sensory cortex, has been a matter of debate (Sreenivasan et al., 2014b; Constantinidis et al., 2018).

Part of the problem in reconciling these contrasting viewpoints is the use of different methodologies across human and animal studies (Miller et al., 2022). In recent years, analysis of intracranial recordings from human subjects have provided evidence for existence of persistent spiking activity during the maintenance of working memory (Kaminski et al., 2017) and other properties of human brain activation during working memory (Haller et al., 2018; Gehrig et al., 2019; Kumar et al., 2021; Xie et al., 2023). Analysis of non-human primate recordings has also begun to mirror methods of analysis inspired by human studies (Rezayat et al., 2021). However differences in behavioral paradigms and methods of analyses persist and make comparison of findings from different models challenging. We were therefore motivated to collect and analyze local field potentials (LFPs) obtained from intracranial recordings in human subjects, using working memory behavioral paradigms and analysis methods that parallel those used in animal models. Our results provide a direct way to compare and reconcile findings in the respective research literatures of the two fields.

## MATERIALS AND METHODS

### Participants

Twelve epilepsy patients participated in the visual working memory study. These were patients with medically intractable epilepsy who had stereo electro-encephalography (sEEG) electrodes implanted to localize their seizure focus. Electrodes were implanted under general anesthesia with implant locations determined on an individual subject basis by clinical considerations related to hypotheses for site of seizure onset. Following surgery, patients were admitted to the epilepsy monitoring unit and antiepileptic medications gradually weaned while patients were continuously monitored for seizures. For our study, we collected LFPs from the contacts of sEEG electrodes using the Natus clinical data acquisition system (Natus, Middleton, WI). All subjects gave informed consent for this study, which was approved by the Institutional Review Board of the Vanderbilt University Medical Center (Nashville, TN, IRB #211037).

### Behavioral Tasks

Participants performed visual working memory tasks using a portable, 13 inch tablet and stylus (Microsoft Surface, Redmond WA) in their hospital rooms, while LFPs were continuously recorded from sEEG electrodes. Task control was achieved in MATLAB 2021 (MathWorks, Natick, MA) using custom Psychophysics Toolbox, Version 3 (Brainard, 1997). At the beginning of each task epoch, the serial port triggers an Arduino Leonardo to send a TTL pulse as timestamps to the trigger channel of the EEG amplifier (Appelhoff and Stenner, 2021).

Variants of a manual delayed response task (spatial task) and a shape match-nonmatch task (shape task) were used. In the spatial task (Fig.1A), a circle appears at the start of each trial in the center of the tablet screen and the subject moves the stylus into the circle to initiate the trial. After a 1 s “fixation” period, a second white circle appears (cue) at a peripheral location for 0.5 s, after which only the center circle remains. This is followed by a 3 s or 6 s (delay) interval where, again, only the fixation point was displayed. After the delay period, the center circle disappears and the subject needs to drag the stylus across the screen into the remembered location of the cue. A TTL pulse marked the beginning of each task event. Delay durations were blocked; trials with 3 s delays were performed first, and then the 6 s delay trials were presented. Cue locations appeared at the periphery of the tablet screen and were evenly distributed across 360 degrees starting at 0 degrees. A typical block had 36 cue locations, each of which appeared once. Cue locations in completed trials were not repeated within a block.

**Figure 1.**
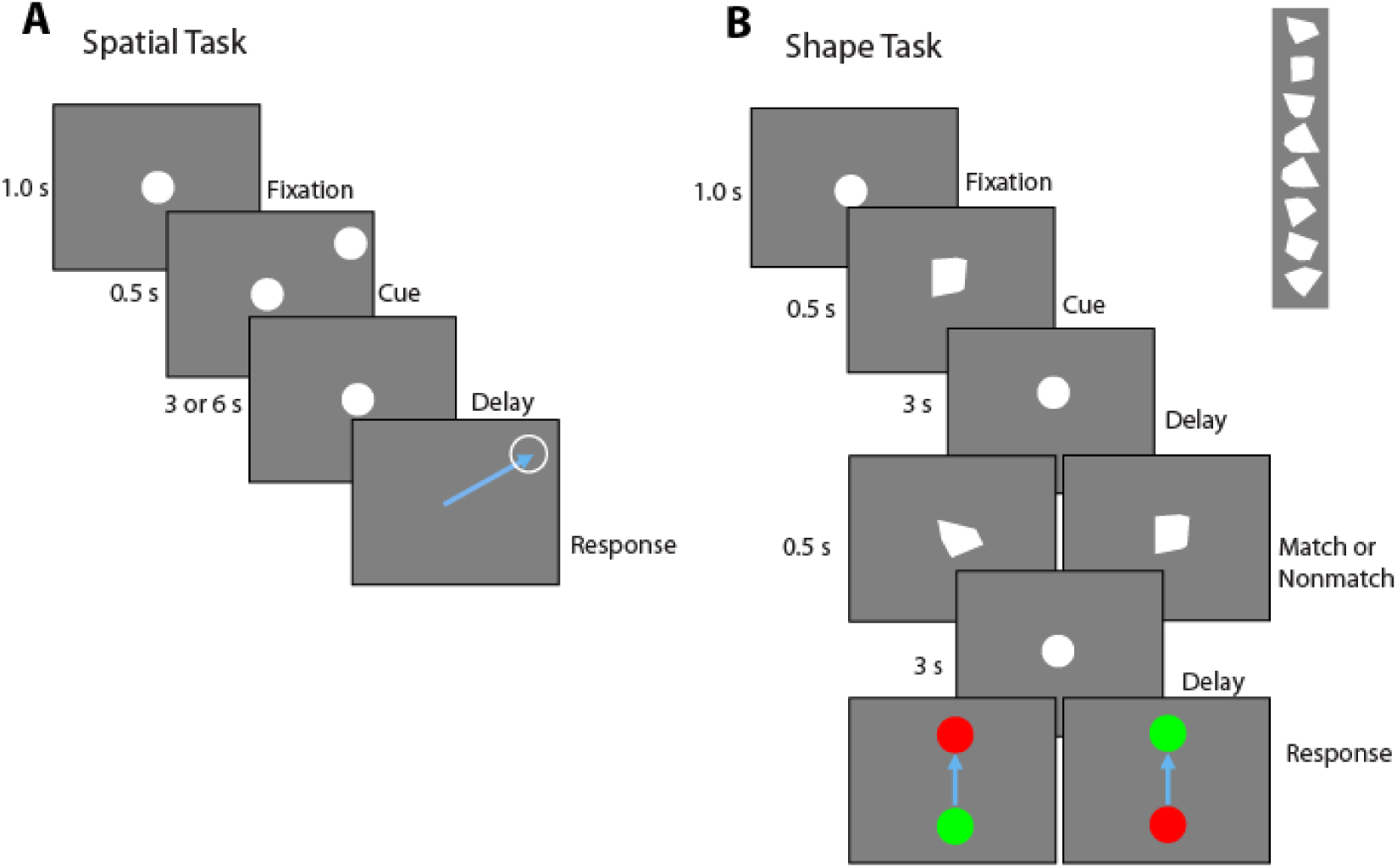
Working memory tasks. A. Spatial Manual Delayed Response Task. At the start of each trial, a circle appears in the center of the tablet screen, and the subject moves the stylus into the circle to initiate the trial. After 1 second, a second white dot appears (Cue) at a peripheral location for 0.5 s, after which only the center circle remains. After a delay period, the center circle disappears and the subject needs to drag the stylus across the screen into the remembered location of the cue. B. Shape Delayed Match to Sample task. At the start of each trial, a white circle appears in the center of the tablet screen, and the subject moves the stylus into the circle to initiate the trial. After a delay period, a white polygon replaces the center circle for 0.5 s (Cue), followed by the reappearance of the center circle. After a delay, a second convex polygon replaces the center circle for 0.5 s, followed by the reappearance of the center circle. After a second delay, the center circle disappears and the subject needs to drag the stylus to either a green or red peripheral circle to indicate whether the two polygons were the same or not.

In the shape task (Figure 1B), at the start of each trial, a circle appears in the center of the tablet screen, and the subject moves the stylus into the circle to initiate the trial. The circle remains visible for a 1 s fixation period. Then a stimulus is presented for 0.5 s, comprising a white, convex polygon (cue), replacing the center circle. This is followed by a 3 s (delay) interval where, again, only the white circle was displayed. A second stimulus (sample) is subsequently shown for another 0.5 s, followed by a second 3 s delay period. At the end of the second delay period, the center circle disappears and two colored circles appeared at the top and bottom of the screen. The subject then needs to drag the stylus across the screen into either the green circle or red circle to indicate whether the two polygons were the same (match) or not (nonmatch), respectively. The polygons used had 3 to 6 vertices, did not extend beyond the extend of the center circle and no polygon was a rotated version of another. The red and green circles’ placement was randomly switched between the top and bottom locations on each trial. A total of eight shapes were used, followed by either a top and bottom location in match or nonmatch to create 32 trials.

Before participants performed the full block of the visual working memory tasks, a member of the experimenter team explained and demonstrated the procedure. The participants then practiced a few trials to ensure they understand the experimental protocol before the block began.

### Electrode Localization

Participants were implanted with multiple electrode shafts of 0.8 mm in diameter and containing 8-16 contacts, spaced between 2.5- 4.3 mm center-to-center (PMT corporation, Chanhassen, MN). We identified the location of all contacts in each individual’s brain by superimposing the pre-surgical MRI and postsurgical CT images using the CRAnial Vault Explorer (CRAVE) software (D’Haese et al., 2012) and the FreeSurfer software package (http://surfer.nmr.mgh. harvard.edu). These software tools allowed us to determine coordinates of each contact, and to identify the anatomical location of each contact according to the Desikan-Killiani Atlas (Desikan et al., 2006) using FreeSurfer’s cortical parcellation and subcortical segmentation procedure (Fischl et al., 2002; Fischl et al., 2004). White matter contacts were excluded from analysis. Electrode contact localizations obtained in this manner were additionally verified by an epileptologist (SWR) based on visual inspection of the co-registered MRI, postsurgical CT and baseline EEG timeseries.

### LFP Recording, preprocessing and signal analysis

Local field potentials were recorded from the implanted sEEG electrodes and sampled at 512 Hz. Task events were synchronized with the LFP recording with a TTL pulse that was generated by the tablet and was recorded in the data acquisition system. Task epochs (e.g. fixation, stimulus presentation, delay period) were aligned to different TTL pulses. LFP recordings were preprocessed by using custom MATLAB code in MATLAB R2022b (MathWorks) and the FieldTrip toolbox (Oostenveld et al., 2011). A bandpass filter between 0.5-200 Hz with a zero-phase sixth-order Butterworth filter were used on single-trial LFP traces. To remove 60 Hz powerline noise and its harmonics, an infinite impulse response (IIR) Butterworth filter was applied. Further, single-trial LFP traces underwent manual inspection for artifact rejection. Electrodes were excluded from data analysis if they were not in gray matter or were determined to be within the patient’s seizure onset zone based on review of seizures by the clinical team and confirmed by a clinical epileptologist (SWR).

The Chronux package (Bokil et al., 2010) was used for time-frequency analysis. We used a multi-taper method to perform a power spectrum analysis of LFPs. The spectrogram of each single trial between 0.5 and 150 Hz was computed with 8 tapers in 500 ms time windows; the spectrograms were estimated with a temporal resolution of 2 ms. We also used the mean filter corresponding to 2 Hz and 2 ms for smoothing the spectrogram of each single trial. In all our analyses, we relied on induced power of the LFP, which is computed by first performing a power computation in each trial and then averaging power across trials. Induced power thus determines power at specific frequencies that may not necessarily be synchronized with specific task events across trials. Power was expressed relative to the mean power recorded during the inter-trial interval. We constructed time-resolved plots (spectrograms) by dividing the power of the signal by the mean inter-trial interval power at each frequency (which is equivalent to subtracting the baseline power in logarithmic, dB, scale). We then standardized the algorithmic power on the temporal profile at each frequency.

### Statistical Analysis

Statistical testing of differences between conditions was performed in the following fashion, using only trials that participants completed correctly. First, we calculated power across an entire task epoch: e.g. fixation period, cue presentation, delay period. Secondly, we averaged power values in these epochs from all available trials of every electrode site, essentially treating each electrode contact as one observation. We then constructed a 1-way or 2-way ANOVA representing the task epoch and region. In every case, the analysis was performed for the beta and high-gamma frequency band, defined as 16-40 Hz and 100-150 Hz, respectively, based on prior studies of working memory (Lundqvist et al., 2016; Haller et al., 2018; Singh et al., 2023). Post-hoc pairwise comparisons on any significant main effects were performed with Tukey’s method.

## RESULTS

### Participants and working memory tasks

Twelve participants were recruited for this study (Table 1). Each participant was implanted with an average of 12.1 ± 2.1 (mean ± SD) sEEG leads containing a total of 154.8 ± 36.3 contacts (Fig. 2), with implant strategy and number of leads determined by clinical considerations. Two stimulus sets were presented (Fig.1A-B). One stimulus set varied the spatial location of a white circle (spatial task), and one involving different polygons shapes (shape task). In the spatial task, the participants had to remember the location of the stimulus presented during the cue period and, after a delay period, indicate where the stimulus appeared by dragging a pointer to the location of the stimulus. In the shape task, two stimuli were presented in sequence with an intervening delay period between them. The subjects needed to determine whether the two stimuli were identical or not and indicate their judgment by selecting a green or red choice target. Participants performed on average 38.4±4.5 trials during the 3s-delay spatial task and 40.9±10.5 trials during the 6s-delay spatial task. Of the 12 total participants, 9 performed 22.4±8.1 trials in the shape task. Data collection lasted approximately 30 minutes, including instructions, practice runs, and pauses between tasks for rest.

**Figure 2.**
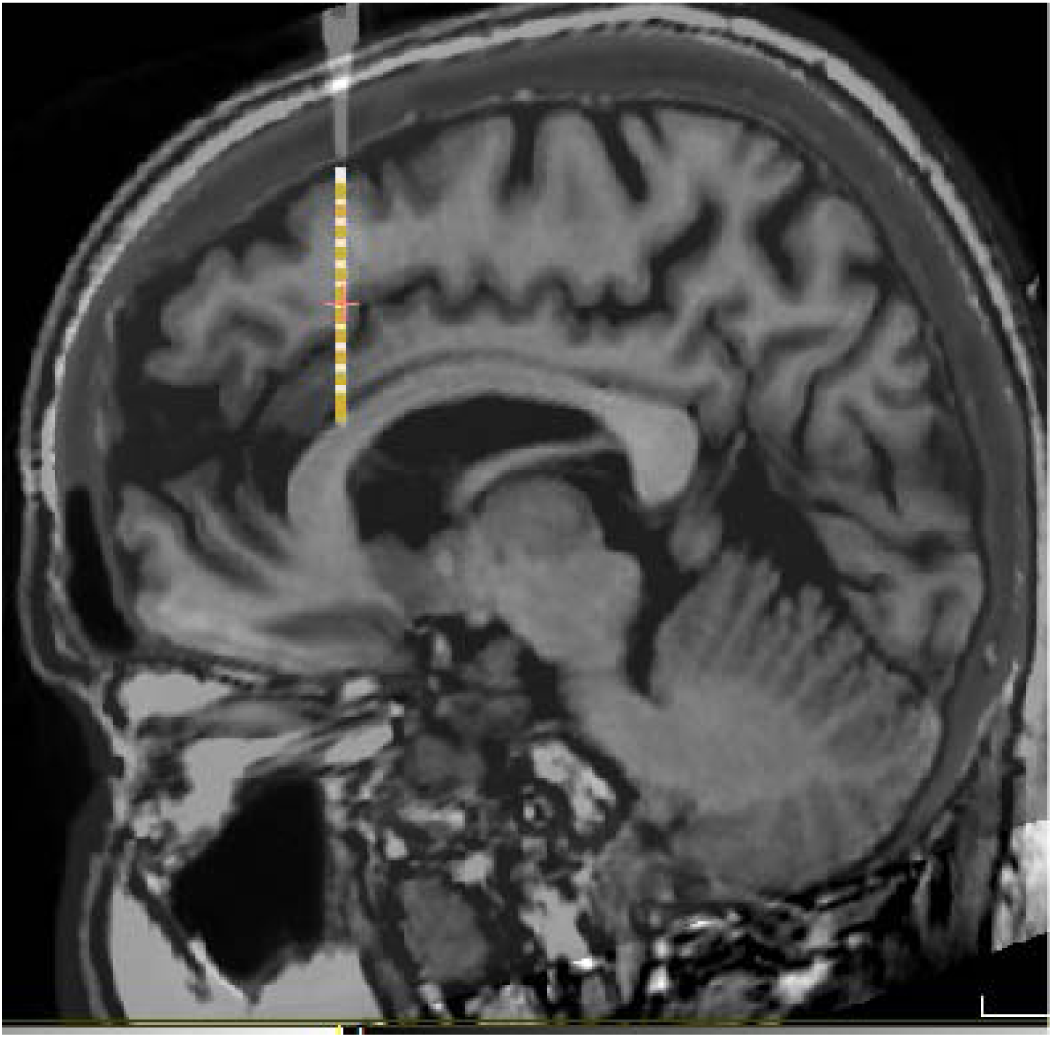
Brain localization. Example MRI scan of one patient, with electrode position, based on CT scan, superimposed, where LFP recording were made.

**Table 1.** Patient demographic characteristics. Table reports age, sex, handedness, and seizure onset, as this was determined clinically, for study participants. Task performance (average across all trials and tasks completed) is also reported.

| Participant Number | Age (years) | Sex | Handedness | Seizure onset | Task Performance |
| --- | --- | --- | --- | --- | --- |
| 1 | 37 | M | Right | Left medial temporal | 90.9% |
| 2 | 22 | F | Right | Left medial temporal<br>Left lateral temporal<br>Left medial frontal<br>Left orbitofrontal | 92.8% |
| 3 | 52 | F | Right | Left medial temporal | 64.0% |
| 4 | 43 | F | Right | Left temporal pole | 42.3% |
| 5 | 29 | F | Right | Left posterior temporal<br>Left occipital | 78.7% |
| 6 | 33 | M | Left | Right medial temporal | 76.8% |
| 7 | 30 | F | Right | Right medial temporal<br>Right lateral temporal | 78.9% |
| 8 | 58 | F | Right | Left medial temporal | 75.0% |
| 9 | 43 | F | Right | Left medial temporal | 74.5% |
| 10 | 33 | F | Right | Right orbitofrontal<br>Right medial temporal<br>Right basal temporal<br>Right parietal | 74.5% |
| 11 | 59 | M | Right | Bilateral medial temporal | 81.3% |
| 12 | 46 | F | Right | Left medial frontal | 86.7% |

### Anatomical localization

LFP data were recorded from cortical and subcortical regions. To provide a concise overview of the patters of LFP activity across the entire brain, we combined recordings into 6 regions, determined based on sample availability, anatomical proximity, and relative similarity of patterns of responses within region. These were defined as follows: a) Mesial Temporal region, which included contacts from the Amygdala, Hippocampus, and Mesial Temporal Cortex (n=11 patients); b) Cingulate region, which included the anterior and posterior cingulate (n=8); c) Lateral Temporal region, which included contacts in the Inferior Temporal Cortex and Superior and Middle Temporal gyrus (n=11); d) Occipital region, which included any contacts in the occipital lobe (n=3); e) Parietal region, which included contacts in the parietal lobe (typically in the posterior parietal cortex, n=7); f) Prefrontal region, which included contacts in any subdivision of the prefrontal cortex (n=10). A few contacts were not localized in any of these regions (e.g. were localized in subcortical structures). These were omitted from analyses presented here. After preprocessing and artifact rejection, signals from a total of 745 contacts from 145 electrode shafts in the 3s-delay spatial task were available (361, 49, 54, 37, 88, 156 contacts in the six regions named above, respectively). Additionally, 724 contacts from 145 electrodes were available in the 6s-delay spatial task (345, 48, 51, 37, 87, 156 contacts in the six regions, respectively). Similarly, 577 contacts from 111 electrodes were available in the shape task (296, 18, 30, 37, 81, 115 contacts in the six regions, respectively). Spectral power was computed in each trial and contact and then averaged within each region.

### Signatures of spectral power during working memory

To examine how oscillatory LFP activity manifested across brain regions and differentiated task events and working memory tasks we plotted the time course of induced LFP power. Mean power in six different brain regions is shown for the two versions (3-and 6s delay) of the spatial working memory task in Fig. 3 and Fig. 4, respectively. Although some differences in the time course of spectral power are present between regions, the more salient finding is the similarity between regions and the engagement of a brain-wide network during the execution of the task. Similarly, mean induced LFP power recorded during the shape working memory task (Fig. 5) revealed brain-wide power modulation during the task.

**Figure 3.**
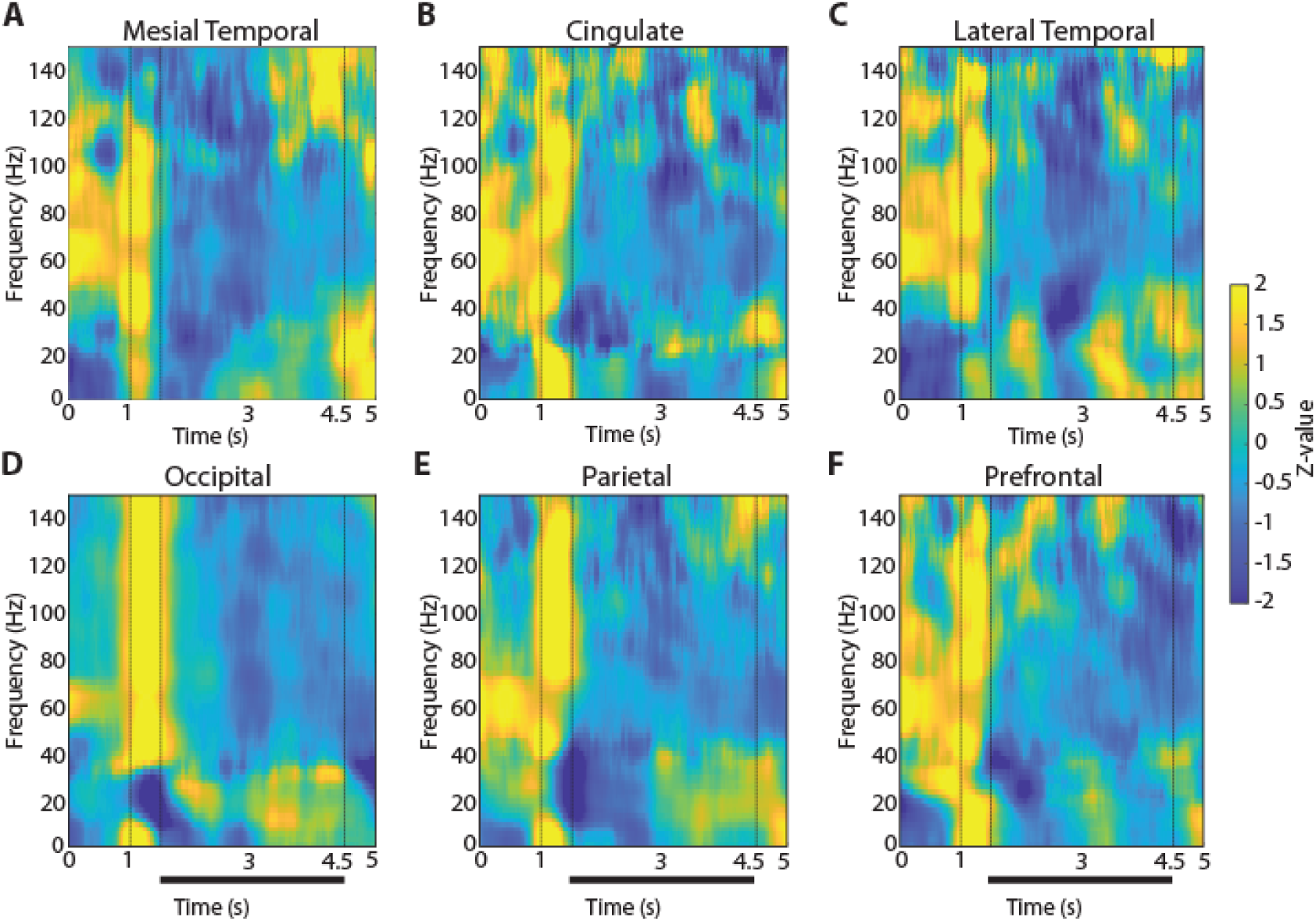
LFP Power spectra across brain regions in the 3-s delay, spatial working memory task. Average, induced power of the local field potential signal relative to baseline (computed in the intertrial interval) is shown for contacts grouped into six brain regions. Vertical lines indicate the time of stimulus presentation (1-1.5 s) and the beginning of the response period (4.5 s). Horizontal bar indicates the delay period of the task over which stimuli needed to be maintained in working memory. **A.** Mesial Temporal regions (Amygdala, Hippocampus, Mesial Temporal Cortex) n= 11 subjects; n=361 electrode contacts, n=6368 trials. **B.** Cingulate regions (Anterior Cingulate; Posterior Cingulate) n= 8 subjects; n=49 contacts, n=1061 trials. **C.** Temporal regions (Inferior Temporal Cortex) n= 11 subjects; n=54 contacts, n=948 trials. **D.** Occipital regions n= 3 subjects; n=37 contacts, n=967 trials. **E.** Parietal regions n= 7 subjects; n=88 contacts, n=2210 trials. **F.** Prefrontal regions n= 10 subjects; n=156 contacts, n=3386 trials.

**Figure 4.**
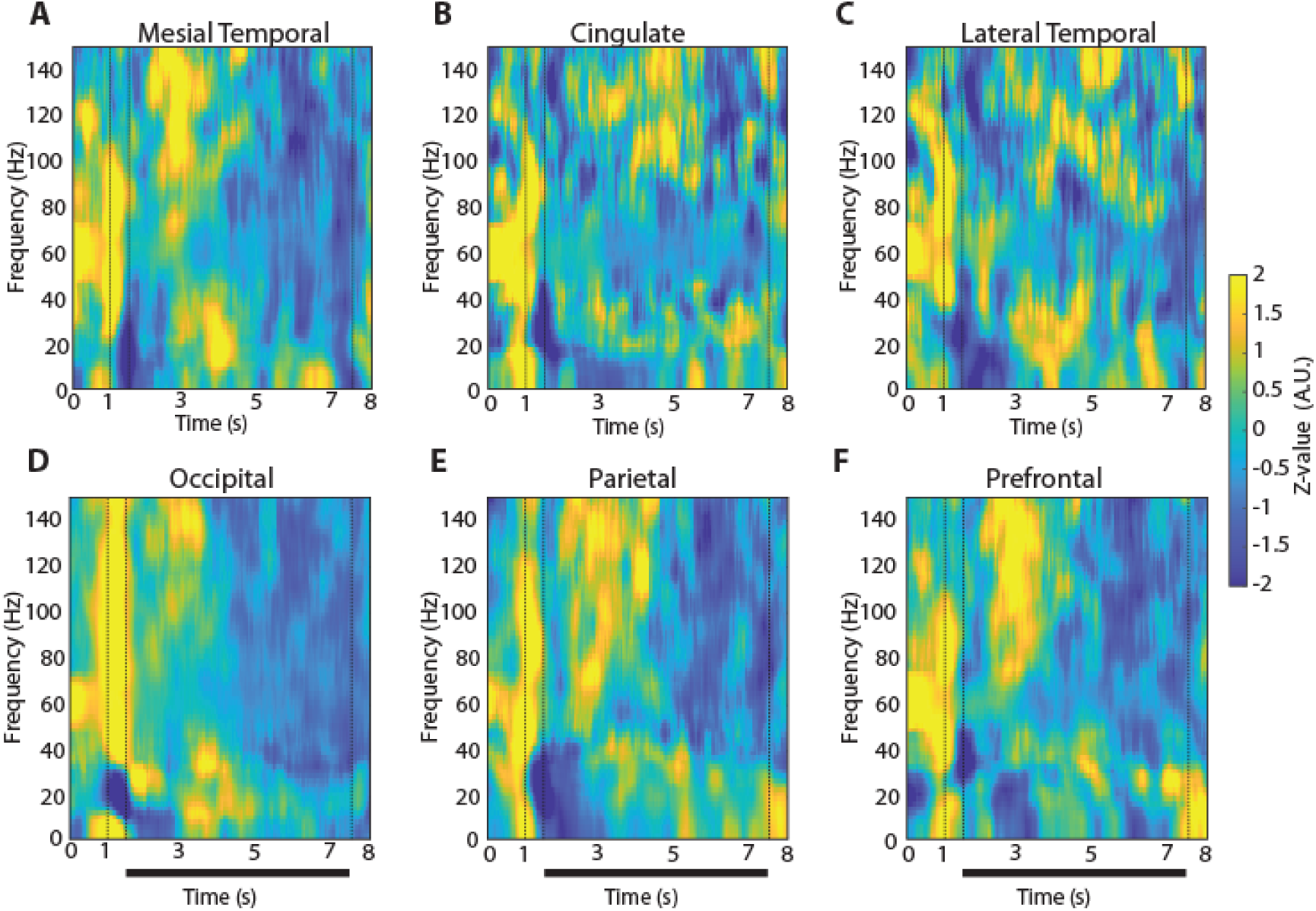
LFP Power spectra across brain regions in the 6-s delay, spatial working memory task. Average, induced power of the local field potential as in Figure 3, now plotted for the 6-s delay version of the spatial working memory task. Vertical lines indicate the time of stimulus presentation (1-1.5 s) and the beginning of the response period (7.5 s). Horizontal bar indicates the delay period of the task over which stimuli needed to be maintained in working memory. **A.** Mesial Temporal regions (Amygdala, Hippocampus, Mesial Temporal Cortex) n= 11 subjects; n=345 electrode contacts, n=5100 trials. **B.** Cingulate regions (Anterior Cingulate; Posterior Cingulate) n= 8 subjects; n=48 contacts, n=953 trials. **C.** Temporal regions (Inferior Temporal Cortex) n= 11 subjects; n=51 contacts, n=737 trials. **D.** Occipital regions n= 3 subjects; n=37 contacts, n=587 trials. **E.** Parietal regions n= 7 subjects; n=87 contacts, n=1570 trials. **F.** Prefrontal regions n= 10 subjects; n=156 contacts, n=2806 trials.

**Figure 5.**
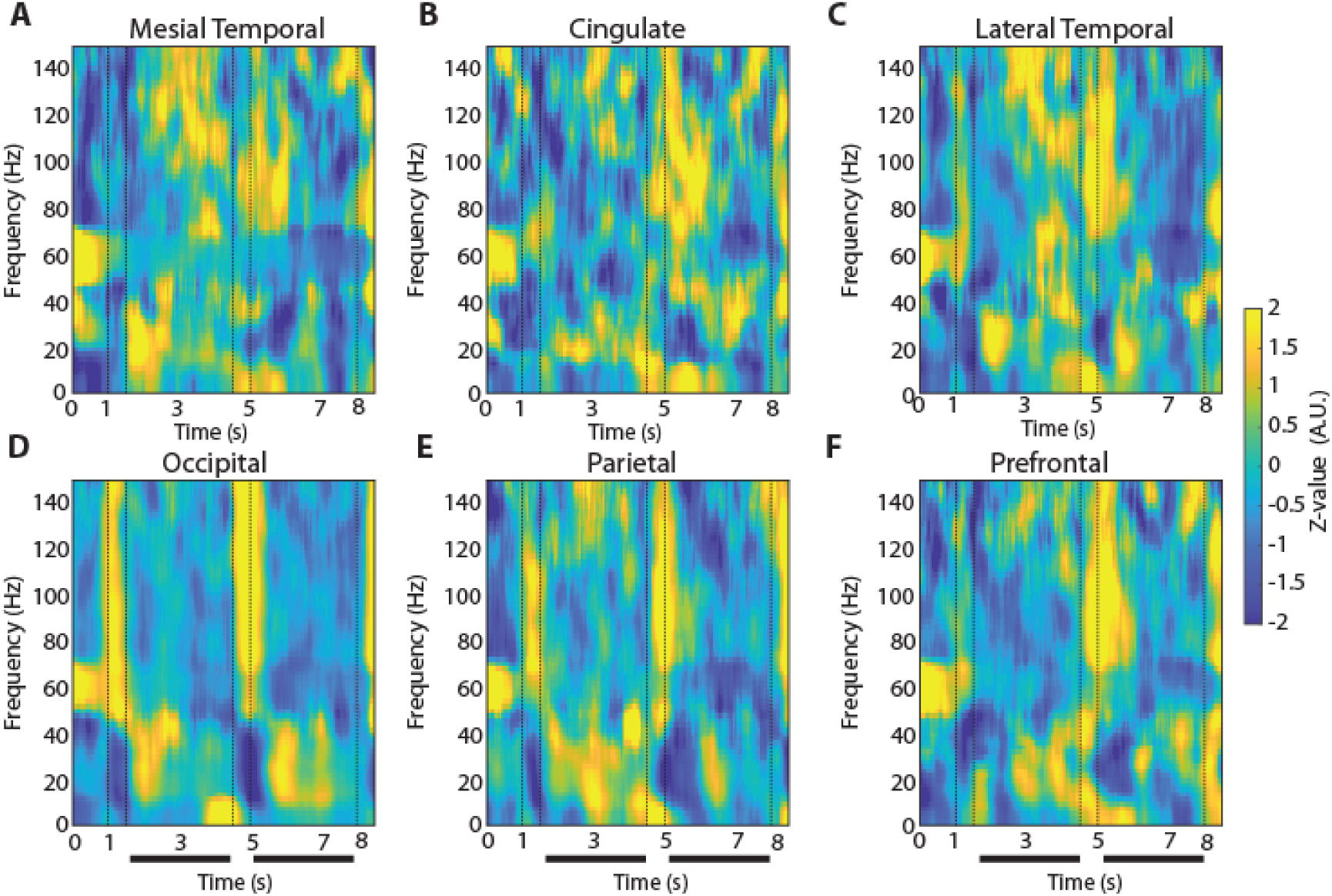
LFP Power spectra across brain regions in the shape, match-nonmatch working memory task. Average, induced power of the local field potential as in Figure 3, now plotted for the shape match-nonmatch, working memory task. Vertical lines indicate the time of the two stimulus presentations (1-1.5 s and 4.5-5) and the beginning of the response period (8 s). Horizontal bars indicate the delay periods of the task over which stimuli needed to be maintained in working memory. **A.** Mesial Temporal regions (Amygdala, Hippocampus, Mesial Temporal Cortex) n= 8 subjects; n=296 electrode contacts, n=5570 trials. **B.** Cingulate regions (Anterior Cingulate; Posterior Cingulate) n= 5 subjects; n=18 contacts, n=329 trials. **C.** Lateral Temporal regions (Inferior Temporal Cortex, Superior and Middle Temporal Gyrus) n= 8 subjects; n=30 contacts, n=522 trials. **D.** Occipital regions n= 3 subjects; n=37 contacts, n=843 trials. **E.** Parietal regions n= 6 subjects; n=81 contacts, n=1912 trials. **F.** Prefrontal regions n= 7 subjects; n=115 contacts, n=2376 trials.

To identify differences between regions and conditions we calculated power in each task epoch, after subtracting at each frequency the baseline power, which was computed in the inter-trial interval. We averaged this power for all trials obtained from each contact in a given electrode and treated it as a single observation (though treating multiple electrode contacts from the same patient, as distinct). The appearance of the cue stimulus in the 3s delay spatial task that participants needed to remember was characterized by broadband increase in power (Fig. 3, vertical lines between 1 and 1.5 s). The most differentiating effect across regions of the cue appearance was the extent of the increase in high-gamma (100-150 Hz) power (1-way ANOVA, F_5,744_ =19.3, p=4.4E-18). The occipital region exhibited the highest power in the high-gamma range compared to all other regions (Tukey post-hoc test, evaluated at the α=0.05 significance level). The full time course of power in the high-gamma band is presented in Fig. 6.

**Figure 6.**
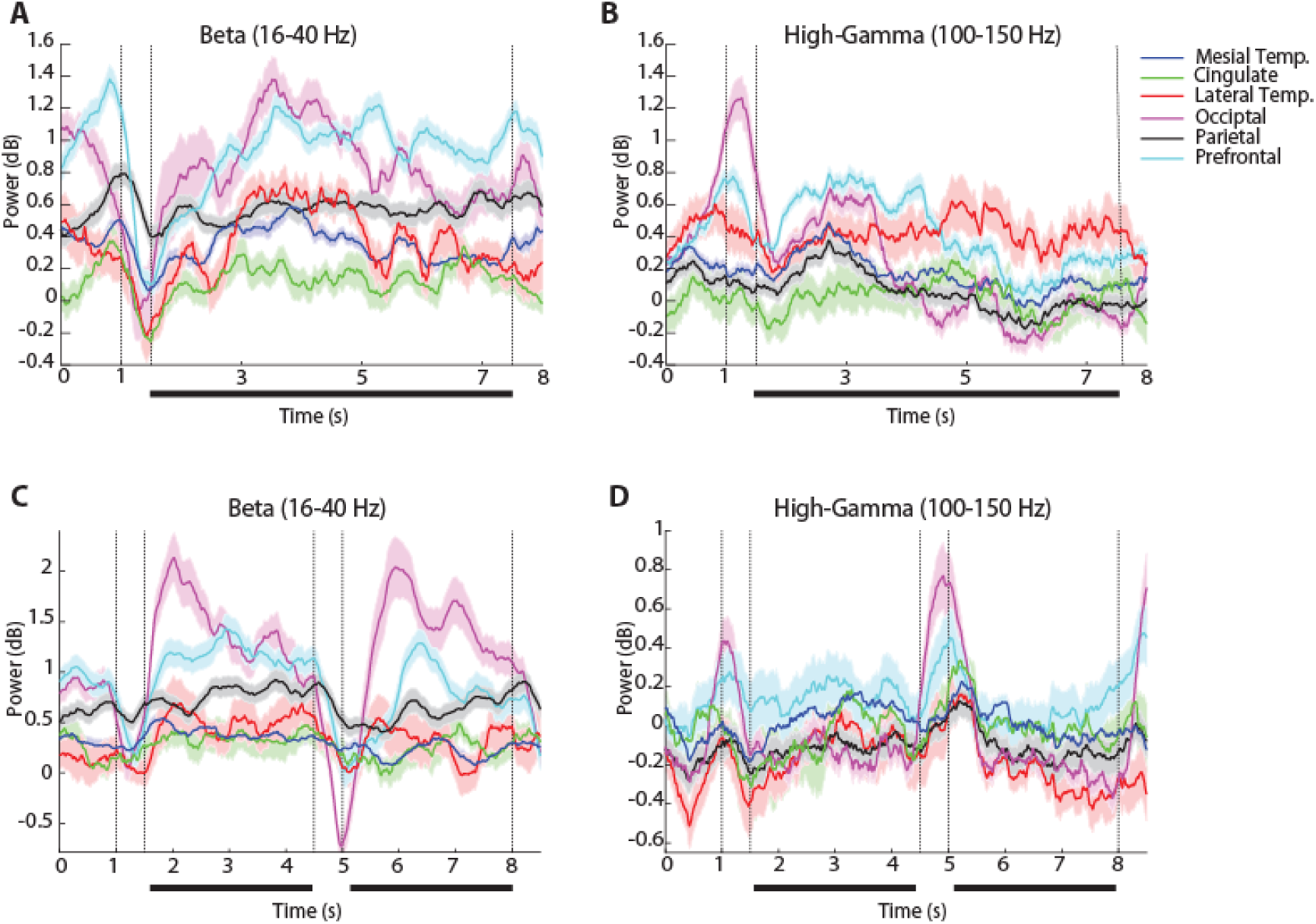
Time course of beta and high-gamma power. **A.** Time resolved induced LFP power in the beta frequency range (16-40 Hz) for the 6-s spatial working memory task, in each of the six brain regions identified. **B.** LFP power in the high-gamma range (100-150 Hz). Similarly, **C.** Time resolved induced LFP power in the beta frequency range (16-40 Hz) for the shape working memory task, in each of the six brain regions identified. **D.** LFP power in the high-gamma range (100-150 Hz).

Another effect of the cue appearance on LFP spectral power in the 3s delay spatial task was a decrease in beta frequency (16-40 Hz). This too differed systematically between regions (1-way ANOVA, F_5,744_ =5.51, p=5.4E-5), with lateral temporal, occipital and parietal regions exhibiting the greatest modulation. The timing of the beta-frequency trough also different between areas. It reached a minimum at the occipital cortex after the stimulus offset, at the parietal cortex 150 ms later, and the prefrontal cortex another 650 ms later. The time course of beta-frequency power can also be seen in Fig. 6.

The patterns of high-gamma and beta power that we identified were generally consistent between versions of the spatial working memory task. In the 6s-delay version of the spatial task, high gamma power again differed significantly between brain regions (1-way ANOVA, F_5,724_ =20.29, p=5.67E-19). The occipital region again exhibited significantly higher power in high gamma frequency compared to all other regions (Tukey post-hoc test, evaluated at the α=0.05 significance level). Similarly, in the beta frequency, there was a significant difference between regions (1-way ANOVA, F_5,724_ =9.15, p=1.9E-08).

The two versions of the spatial task imposed different difficulty in terms of their working memory requirement. We examined how this translated into spectral power across regions, focusing again in the high-gamma range (Figure 7). We adopted a 2-way ANOVA with factors regions, and 3s/6s version of the spatial task. Here we considered the first 3 s of the delay period in the two task versions to compare power. High gamma power was higher in the 6s-delay version of the task (F_1,1469_ =50.6, p=1.8E-12 for main effect of task). We also found an interaction between regions and versions of the spatial task, although this was at the margin of statistical significance (F_5,1469_ =2.38, p=0.04). The parietal region (Fig. 7) exhibited significantly higher power compared to other regions (F_5,1469_ =6.44, p=6.3E-06, Tukey post-hoc test, evaluated at the α=0.05 significance level). It was also notable that the elevated high-gamma power the 6s version of the task was already present in the fixation period, prior to the appearance of the stimulus or the delay period. Since the tasks were presented in blocks, the result suggests that the participants’ level of effort or expectations about the task modulated systematically spectral power.

**Figure 7.**
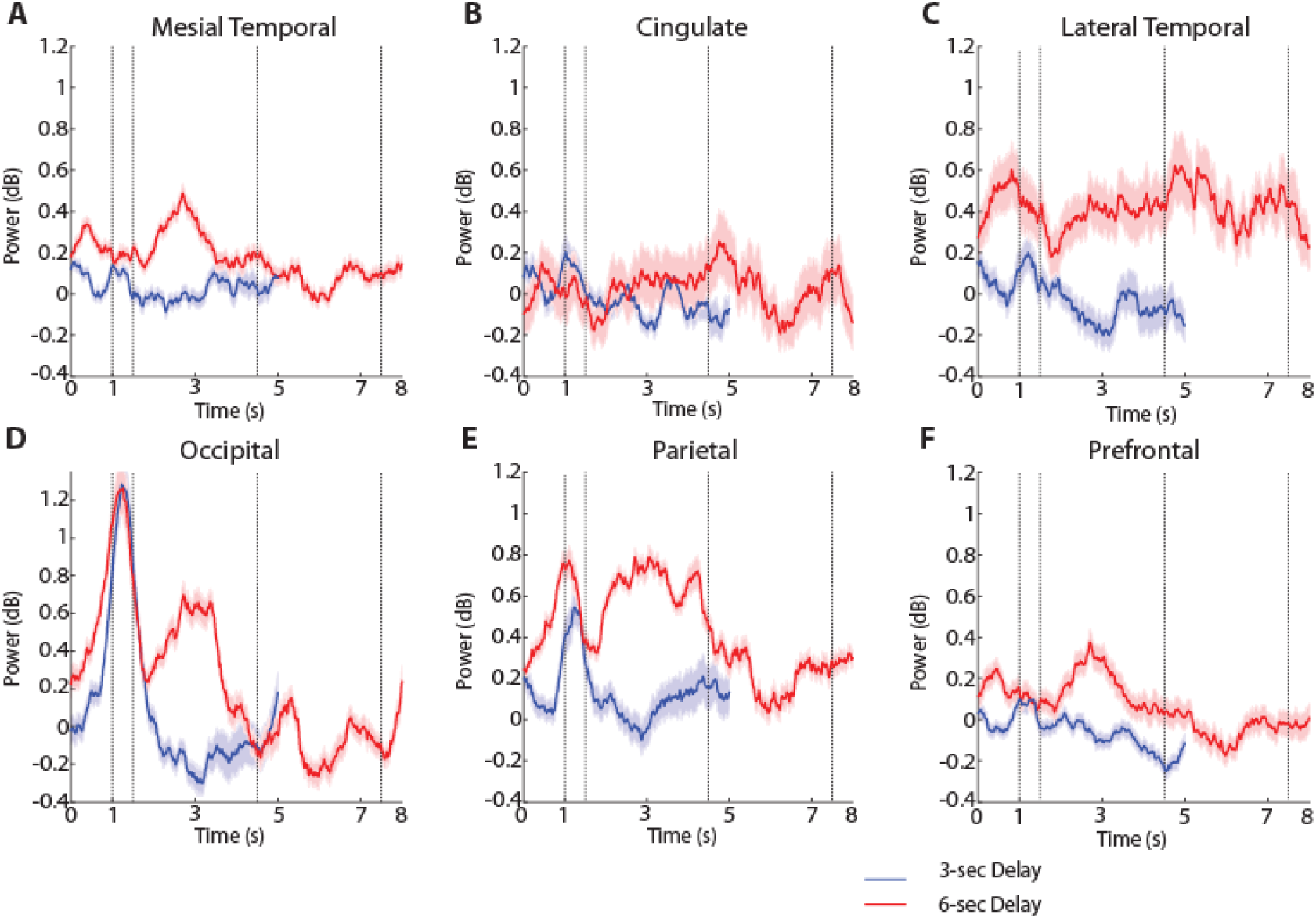
High-gamma power for different delay durations. Time resolved induced LFP power in the high-gamma range (100-150 Hz) for the two versions of the spatial working memory task, involving 3 s and 6 s delay periods. **A.** Mesial Temporal regions. **B.** Cingulate regions. **C.** Temporal regions. **D.** Occipital regions. **E.** Parietal Regions. **F.** Prefrontal regions.

### Spatial vs object working memory

LFP power was also analyzed in the shape task (Fig. 1B), which differed in the nature of stimuli that needed to be maintained in memory (shapes rather than spatial locations), the part of the visual field where the stimuli appeared (central, vs. peripheral) but also in the nature of the task (requiring a judgment on whether the second stimulus was match or nonmatch and preparation of a motor response based on a color rule). High gamma power during the cue presentation period differed systematically between regions, for this task, too (1-way ANOVA, F_5,576_ =3.18, p=0.0077). Beta-frequency power during the cue presentation also differed between regions (1-way ANOVA, F_5,576_ =4.57, p=0.0004). However, the time course of beta power was distinctly different than what we observed in the spatial task (Fig. 6): a brief decrease in beta power during the cue presentation was followed by a rebound above the baseline, and then a second decrease around the time of the second stimulus presentation.

To perform the task, it was essential for subjects to maintain the identity of the stimulus in memory during the first delay period. Thus, we examined how the gamma differed between regions as the subjects were engaged in this function (Fig. 5, 6). Unexpectedly, we found high gamma power was not significantly different between regions (1-way ANOVA, F_5,576_ =1.76, p=0.12), though the prefrontal cortex was the area with the highest power throughout the delay period of the task (Fig. 6D). Instead, beta power differed systematically between regions during the first delay period, (1-way ANOVA, F_5,576_ =13.46, p=2.0E-12), with the occipital region exhibiting the greatest amplitude (Fig. 6C).

## DISCUSSION

Our study revealed patterns of oscillatory brain activity during the execution of visual working memory tasks including widespread and region-specific patterns of activity visible in the LFP. The appearance of a visual stimulus across all tasks generated broad-band increases in LFP activity. Regions were differentiated by subtle but consistent signatures of activity in the high-gamma and beta frequency ranges. High-gamma activity associated with stimulus presentation was most pronounced in the occipital (visual) cortex and beta activity was suppressed with a region-specific time course from occipital to parietal to prefrontal and temporal areas. These results are consistent with hierarchical processing of visual information and support the idea that memory may travel along the sensory processing stream of the utilized sensory modality (Kucewicz et al., 2019).

### Regional localization of working memory

The locus of working memory maintenance in the brain has been a matter of debate in recent years. Neurons in the prefrontal cortex and areas connected with it generate persistent discharges tuned for stimulus properties during the delay intervals of working memory tasks, in animal (Funahashi et al., 1989), and human intracranial recordings (Kaminski et al., 2017). Computational models typically simulate persistent activity in neural networks with recurrent connections between units with similar tuning for stimuli (Compte et al., 2000), which capture working memory behavior very well, particularly in spatial working memory tasks (Wimmer et al., 2014). Based on these results, it has been postulated that the prefrontal cortex plays the primary role in the maintenance of working memory by virtue of generation of persistent spiking activity (Constantinidis et al., 2018).

However, this idea has been challenged. Human imaging studies have applied Multi-Variate Pattern Analysis (MVPA), examining the simultaneous pattern of activation of multiple voxels to different task conditions (Offen et al., 2009), in order to successfully decode working memory content from the primary (Harrison and Tong, 2009; Albers et al., 2013; Xing et al., 2013) and extrastriate visual cortex (Tong and Pratte, 2012; Ester et al., 2013; Sreenivasan et al., 2014a). On these grounds, it has been suggested that the prefrontal cortex may play a supervisory or control role in working memory, “highlighting” the locations of stimuli held in memory while the contents of working memory are maintained in sensory cortex (Sreenivasan et al., 2014b).

This is not to say, however, that models of prefrontal persistent activity are inconsistent with our findings. It is understood that a distributed network of cortical and subcortical areas generates persistent activity during working memory (Constantinidis and Procyk, 2004; Christophel et al., 2017). The prefrontal cortex is essential for the ability of the network to maintain persistent discharges, by virtue of the biophysical properties of its neurons and pattern of connectivity (Leavitt et al., 2017; Mejias and Wang, 2022). In our results, high gamma power was consistently elevated in the prefrontal cortex during the delay period across tasks (Fig. 6B, D), although this difference was quantitative rather than qualitative in comparison to other areas, and gamma power may be an imprecise index of working memory (as discussed in the next section). The presence of oscillatory processes in other brain areas was also revealing. Mesial temporal structures, such as the hippocampus and amygdala, play well understood roles in long-term memory (Burgess et al., 2002; Moscovitch et al., 2016). More recent studies, however, suggest engagement during working memory processes, as well (Kaminski et al., 2017; Borders et al., 2022). Our results confirmed these findings and suggested modulation of oscillatory processes by the working memory tasks.

### Basis of working memory in neural activity

As discussed, models of working memory informed mostly by animal studies have identified persistent discharges generated in the prefrontal cortex and other areas as the critical neural correlate of working memory (Constantinidis et al., 2018). Such persistent spiking generation is most often associated with gamma frequency oscillations in the LFP (Pesaran et al., 2002). Some recent working memory models have emphasized the rhythmicity of spiking discharges themselves and posited that bursting in the gamma frequency range is the critical variable that tracks stimulus information maintenance in working memory (Lundqvist et al., 2016; Miller et al., 2018). Human intracranial recordings, too, reveal rhythmicity in the gamma band during working memory tasks (Tallon-Baudry et al., 1998; Tallon-Baudry et al., 2001; Haller et al., 2018). Critical task parameters, such as working memory load, have been shown to modulate neural oscillations (Jensen et al., 2007; Roux and Uhlhaas, 2014). For this reason, gamma power is considered a marker of task-related activation (Crone et al., 2006).

However, gamma frequency oscillations in the LFP is at best an imprecise index of neural activity mediating working memory. The LFP represents summation of ionic currents in a cortical volume, in the order of 0.1-0.2 mm, and are driven by both spiking and synaptic events, e.g. postsynaptic potentials propagated from distant areas that fail to generate action potentials in the area where the recording takes place (Buzsaki, 2004; Kajikawa and Schroeder, 2011). During presentation of stimuli, correlated bottom-up inputs can serve to synchronize population neuronal spiking, and phases of synchronized excitation by pyramidal neurons followed by inhibition by interneurons can thus produce rhythmicity specifically in the gamma frequency range (Fries, 2009). Less precisely-timed or correlated inputs may fail to generate gamma oscillations, and indeed recent animal studies have suggested more prominent changes in the beta rather than gamma frequency range after learning to perform working memory tasks (Singh et al., 2022; Singh et al., 2023). Similarly, a recent human study of activation patterns in auditory working memory with intracranial recordings demonstrated that frontal and temporal regions with high decoding accuracy were not accompanied by significant increases in gamma power (Uluç et al., 2023).

Beta frequency oscillations are thought to represent a signature of activity during top-down processes, that is disrupted by appearance of exogenous, sensory stimuli (Haegens et al., 2011; van Kerkoerle et al., 2014; Bastos et al., 2015). Beta oscillations are readily detectible in other extracellular field recordings (such as EEG or MEG) and are also a reliable marker of underlying cognitive processes impacting neural circuit interactions, just as gamma oscillations are (Uhlhaas and Singer, 2011; Roux and Uhlhaas, 2014), including working memory and top-down control (Siegel et al., 2012; Helfrich and Knight, 2016). Consistent with the aforementioned animal studies (Singh et al., 2022; Singh et al., 2023), in our study, decrement of beta oscillations was detected during the task execution, and differed systematically between areas, at least around the time of stimulus presentations and early in the delay period of the task.

### Study limitations and open questions

Although our analysis focused on trials completed correctly, it is important to note that epilepsy patients are known to suffer from cognitive deficiencies, including in frontal lobe function and working memory (Stretton and Thompson, 2012). In this respect, patterns of brain activity we describe here may deviate systematically from those of healthy participants.

Additionally, our results relied entirely on LFPs. Although providing an informative reflection of neural activity, LFP power does not map strictly onto spiking activity (Buzsaki, 2004). Results from combined methodologies, including neuron recordings from large populations of isolated neurons in humans and animal models, and combined single-neuron and LFP analysis in the near future are making it possible to study in detail the role of areas and patterns of neuronal activity in working memory.

## Acknowledgments

Research reported in this paper was supported by NIH grants R01 EY017077 (CC) K23 AG072030 (SWR) and K12 NS080223 (SKB). We wish to thank Janki Bava, Rye Jaffe and Junda Zhu for helpful comments on the manuscript.

## Author Contributions

Conducted research: BS, ZW, LMM, EG, JNF, DJE, SKB, SWR, CC. Analyzed data: BS, ZW, GWJ, SWR, CC. Wrote the manuscript with input from all authors: BS, CC.

